# A pseudovirus-based method to dynamically mimic SARS-CoV-2-associated cell-to-cell fusion and transmission

**DOI:** 10.1101/2023.06.26.546514

**Authors:** Xiangpeng Sheng, Yi Yang, Fang Zhu, Fan Yang, Honghua Wang, Ronggui Hu

## Abstract

SARS-CoV-2 has caused the global tremendous loss and continues to evolve to generate variants. Entry of SARS-CoV-2 into the host cells is primarily mediated by Spike (S), which binds to the host receptor hACE2 and initiates virus-cell membrane fusion. Cell fusion contributes to viral entry, cell-to-cell transmission and tissue damage in COVID-19 patients. Many reporter assays have been developed to study S-mediated cell fusion by equally coculturing S-expressing cells and hACE2-positive cells. However, these strategies cannot fully simulate cell-to-cell fusion and transmission of SARS-CoV-2 infection, in which virions from a single target cell transmit to the neighbor cells and induce syncytia formation. Here, we design a pseudovirus-based method to dynamically mimic cell-to-cell fusion and transmission of SARS-CoV-2. We coculture a small number of pseudovirus-producing 293FT cells and a large number of hACE2-expressing 293T cells, and demonstrate that a single cell producing S-pseudotyped virions can induce significant syncytia of hACE2-positive cells. This pseudovirus-based method is a powerful tool to screen and estimate potential inhibitors of S-driven syncytia. Moreover, this strategy can also be utilized to explore fusogenic ability of SARS-CoV-2 variants. Together, the pseudovirus-based method we report here will be beneficial to drug screening and scientific research against SARS-CoV-2 or future emerging coronavirus.

## Main contents

Severe acute respiratory syndrome coronavirus 2 (SARS-CoV-2), responsible for the COVID-19 pandemic, has caused the global tremendous loss and continues to evolve to generate variants. Entry of SARS-CoV-2 into target host cells is primarily mediated by spike (S), which binds to the host receptor hACE2 and initiates virus-cell membrane fusion [1]. Most COVID-19 patients had showed pneumocyte syncytia in the lungs [2]. Cell fusion contributes to viral entry, cell-to-cell transmission and tissue damages, and thus attracts much attention. Because authentic SARS-CoV-2 live virions can only be handled in the Biosafety Level 3 (BSL-3) facilities, many researchers have developed different assays to study cell fusion in BSL-1/2 by directly expressing S and hACE2 on mammalian cells [2-5]. Briefly, in the regular cell fusion assays, S-expressing cells and hACE2-positive cells are cocultured at about a 1:1 ratio, which induces cell-cell fusion and usually activates a fusion reporter. Although these strategies are useful, they cannot efficiently simulate cell-cell fusion and transmission in SARS-CoV-2 infection, in which virions from one target cell transmit to neighbor cells and result in syncytia. Here, we design a pseudovirus-based method to dynamically and highly mimic cell-to-cell fusion and virus transmission of SARS-CoV-2.

First of all, we generated spike-pseudotyped virions (S pseudovirions) in HEK293FT cells by co-transfecting three plasmids, including psPAX2, pCDH-sfGFP, and a plasmid expressing SARS-CoV-2 S (Figure 1A), and collected the viral supernatant, which is similar to a previous report [6]. As shown, S pseudovirus efficiently infected hACE2-positive 293T cells (293T-hACE2), but not the control 293T or 293FT cell lines (Supplementary Figure S1A, B), suggesting that S indeed envelops the pseudovirions. However, fluorescent microscopy revealed that the infection of 293T-hACE2 cells by pseudovirus supernatant could not trigger the cell-cell membrane fusion events (Supplementary Figure S1C). Given that the authentic SARS-CoV-2-infected host cells can continue to generate live virions to promote formation of the syncytia, we hypothesized that a cell producing pseudovirions may show a similar capacity to induce cell-cell fusion.

**Figure 1.**
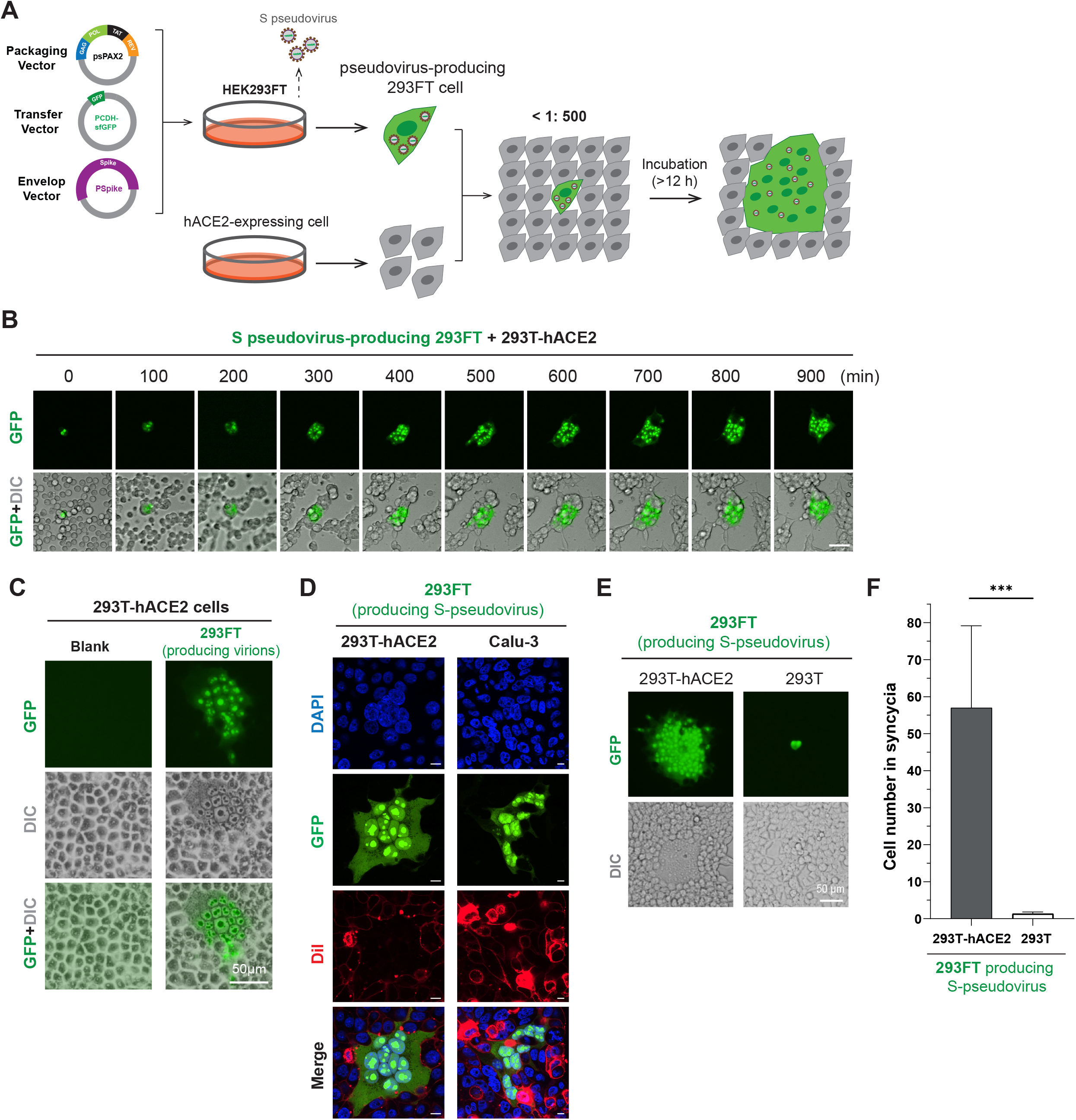
A novel strategy to mimic cell-cell fusion and transmission of SARS-CoV-2. **(A)** The schematic illustration of pseudovirus-based cell-cell fusion assay. In Brief, psPAX2, pCDH-sfGFP and an S-expressing plasmid were co-transfected to HEK293FT cells for 24 h. Pseudovirus-producing 293FT and hACE2-expressing 293T cells were cocultured at a ratio of about 1:500. Syncytia were monitored under a real-time imaging system or imaged at indicated times. **(B)** Real-time imaging of syncytia formation. Pseudovirus-producing 293FT and 293T-hACE2 cells were mixed and seeded in 29 mm dishes, and monitored by the Olympus SpinSR real-time live cell imaging system. Scale, 50 μm. **(C)** The syncytia had a large cytoplasm containing multiple nuclei. The images were taken after 15 h coculture of pseudovirus-producing 293FT and 293T-hACE2 cells, using Olympus IX73. Scale, 50 μm. **(D)** Confocal imaging of syncytia with nuclear and plasma membrane staining. Syncytia of 293FT (left) or Calu-3 (right) cells were fixed by 4% PFA, and then subjected to membrane staining (10 μM Dil, 30 min) and nuclear staining (DAPI). Scale, 10 μm. (**E** and **F**) Pseudovirus-producing 293FT cocultured with hACE2-negative 293T cannot induce cell fusion. Virions-producing 293FT cells were mixed with 293T or 293T-hACE2 for 24 hours. (E) shows the microscope images, and (F) is the quantitative analysis of the relative sizes of syncytia. Scale, 50 μm. Data are present as mean ± SD, \*\*\**P* < 0.001. (Student’s *t*-test, n > 10).

To test the hypothesis, pseudovirus-producing 293FT cells (GFP-positive) were digested to be single cells after 24 h transfection, and were subsequently cocultured with 293T-hACE2 cells at a 1:500 ratio (Figure 1A). The ratio of 293T-hACE2 cells to pseudovirus-producing 293FT cells in the mixture should be very high (about 500:1), otherwise the formed syncytia may not be derived from a single pseudovirus-producing 293FT [5]. Real-time live cell imaging revealed that a single 293FT cell producing pseudovirions can readily induce a large syncytia formation in 293T-hACE2 (Figure 1B and Supplementary Video S1). The virions-producing GFP-positive 293FT can fuse with the closest neighbor cells first, which facilitates transmission of the pseudovirions to the neighbor cells and further promotes more and more cell-cell fusion. Moreover, the size of the GFP-positive syncytia is dependent on the coculture time (Figure 1B). In the syncytia, multiple and separate nuclei were clearly observed to pack together (Figure 1C). Confocal imaging of cells with nuclear and membrane staining further confirmed the membrane fusion of the syncytia (Figure 1D). In addition, pseudovirus-producing 293FT cells can also trigger syncytia formation in Calu-3 cell line, a human bronchial epithelial cell line that endogenously expresses hACE2 (Figure 1D), which suggests that any hACE2-expressing cells cocultured with virions-producing cells could form syncytia. Notably, different from hACE2-expressing 293T, regular 293T cells cocultured with pseudovirus-producing 293FT cannot cause syncytia formation (Figure 1E, F), confirming that the syncytia were S/hACE2-driven cell-cell fusion. The data also excluded the possibility that the division or proliferation of the virions-producing cell might contribute to syncytia formation. Therefore, a single cell producing S-pseudotyped virions can induce syncytia of hACE2-positive cells, which highly mimic authentic SARS-CoV-2-induced cell-cell fusion and transmission.

In a previous study, several most effective drugs, including niclosamide and salinomycin, had been identified to inhibit syncytia expansion; Non-competitive SERCA inhibitors thapsigargin (TG) and cyclopiazonic acid (CPA) causing Ca^2+^ depletion also show an inhibitory effect on syncytia formation [2]. Our pseudovirus-based method was therefore utilized to test these potential drugs. Consistently, niclosamide, salinomycin and TG dramatically block the formation of syncytia mediated by wild-type (WT) S-pseudovirus-producing 293FT (Figure 2A, B). CPA treatment also disrupted virions-mediated cell fusion, but its inhibitory function was much weaker than other drugs. Furthermore, this phenotype was recapitulated by Delta S-pseudovirus-producing cells (Figure S1D and Supplementary Figure S1E). Therefore, the pseudovirus-based strategy can be used to estimate the inhibitory effects of potential compounds that may restrict cell fusion, which makes it a powerful tool for drug screening.

**Figure 2.**
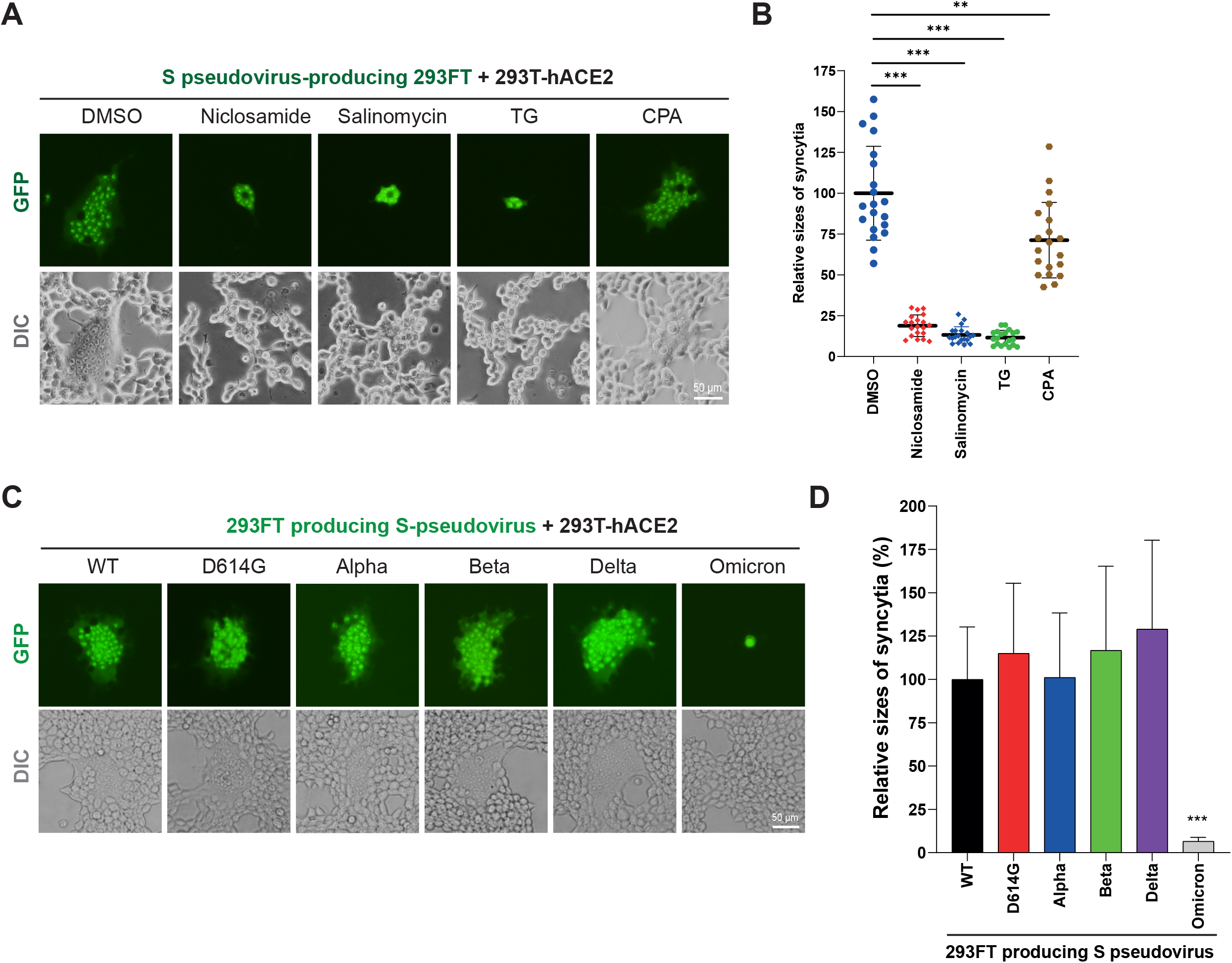
This pseudovirus-based strategy is a powerful tool for drug screen and SARS-CoV-2 investigation. **(A)** This pseudovirus-based method is a powerful tool to estimate the inhibitory effects of potential drugs on S-mediated cell-cell fusion. DMSO, Niclosamide (1μM), Salinomycin (1μM), thapsigargin (TG) (1μM) or cyclopiazonic acid (CPA) (5μM) was added to the coculture system of pseudovirus-producing 293FT and 293T-hACE2 cells. Scale, 50 μm. **(B)** Quantitative analysis of the syncytia sizes in (A). Data are shown as mean ± SD, \*\**P* < 0.01, \*\*\**P* < 0.001. (Student’s *t*-test, n = 20). **(C)** This pseudovirus-based strategy revealed that Omicron had less fusogenic activity than early-pandemic SARS-CoV-2 variants. 293FT cells producing different S pseudovirions were separately cocultured with 293T-hACE2 for 24 h, and syncytia were imaged by Olympus IX73. Scale, 50 μm. **(D)** Quantitative analysis of the syncytia sizes in (C). Data are present as mean ± SD, \*\*\**P* < 0.001. (Student’s *t*-test, n = 10).

SARS-CoV-2 evolutes to generate many variants of concern (VOCs) having increased infectious abilities and changed pathogenicity [7]. Particularly, Omicron had been reported to be less fusogenic than early-pandemic SARS-CoV-2 variants [8, 9]. We constructed 293FT cells generating different pseudovirions carrying distinct S variants, and conducted the cell-cell fusion assay. As shown, while most S variants can induce significant syncytia formation, Omicron showed a very weak ability to mediate cell-cell fusion (Figure 2C, D), consistent to the previous reports. Thus, this method shows the potential to explore the fusogenic capacity of the SARS-CoV-2 variants.

Taken together, we develop a pseudovirus-based method to dynamically investigate cell fusion and cell-to-cell transmission of SARS-CoV-2, which will be useful for the researchers who do not have access to a BSL-3 facility. This method could be used for high-throughput screening of drugs that have abilities to inhibit S mutants-induced syncytia formation and virus transmission. However, to provide sufficient area for the formation of at least 3 syncytia in each well, 48- or 96-well plates, but not 384-well plates, should be used in a high-throughput screening assay, which may increase reagent costs and screening time; moreover, it may be a bit difficult to ensure same syncytia numbers in each well. Importantly, given that other coronaviruses may have similar features to result in cell-cell fusion and syncytia formation in host, we believed that this strategy we design here will be very beneficial to scientific research of SARS-CoV-2 variants or future emerging coronavirus.

## Materials and Methods

### Cell lines

Human embryonic kidney cell line 293T and 293FT, and Calu-3 cells were cultured in Dulbecco’s Modified Eagle Medium (DMEM) containing 100 units/ml penicillin, 100 mg/ml streptomycin, 2 mM L-glutamine and 10% fetal bovine serum (FBS), at 37 °C with 5% CO_2_. 239T cell line stably expressing human ACE2 (293T-hACE2) was constructed by hACE2 overexpression.

### Plasmids and reagents

pSPAX2 was obtained from Addgene. pCDH-sfGFP was provided by Dr. Qiang Deng from Fudan University. The pVAX1 vectors encoding several S variants (WT, D614G, Alpha, Beta, Delta) were generously offered by Prof. Dimitri Lavillette from Institut Pasteur of Shanghai. Omicron S gene was synthesized and cloned to pVAX1.

The anti-GFP (#RLI-09) mouse antibody was provided by Biolinkedin, Shanghai. Niclosamide (HY-B0497) and Salinomycin (HY-15597) were ordered from MCE. Thapsigargin (SC0389-2mM) was obtained from Beyotime Biotechnology, and cyclopiazonic acid (T15027) was from TargetMol Chemicals Inc.

### Production and transduction of SARS-CoV-2 S pseudovirions

Similar to a previous research [6], psPAX2, pCDH-sfGFP and a S-expressing plasmid (ratio: 4.5:3:3) were co-transfected into 293FT cells with Lipofectamine™ 2000 (Invitrogene). 48 h later, virus supernatants were collected and passed through 0.45 μm filters, and then stored at -80 °C. For pseudovirus transduction, 293T-hACE2 cells were digested to be single cells by trypsin-EDTA buffer. 293T-hACE2 cells were incubated in 1 mL DMEM containing pseudovirus, and seeded in 12-well plates at 37 °C. After 24 h, culture media were changed to fresh DMEM. Cells were further cultured for 24 h, and digested by trypsin. Flow cytometric analysis was used to determine the percentages of GFP-positive cells.

### Pseudovirus-based cell-cell fusion assay

To prepare virions-producing cells, psPAX2, pCDH-sfGFP and a S-expressing plasmid were co-transfected into 293FT cells at a ratio of 4.5:3:3 with Lipofectamine 2000 for 24 h. Pseudovirus-producing 293FT cells and 293T-hACE2 cells were detached into single cells by using trypsin, and cocultured at a ratio of more than 1:500. Cell mixture was then maintained in 37 °C for 12∼24 hours. Live cell imaging was conducted by Olympus IX73 or Olympus SpinSR system.

### Confocal fluorescence microscopy

According to a previous confocal protocol [10], cells were fixed in 4% PFA at room temperature for 20 min, and washed with DPBS for three times. Cells were then stained with 10 μM DiI (red) for about 30 min. After washing cells with DPBS, DAPI was utilized for nuclear staining. Confocal imaging was performed by Olympus SpinSR imaging system.

### Statistical analysis

All data are present as mean ± SD. GraphPad Prism was used for data analysis. Statistical significance was determined by Student’s *t*-test. *P* values < 0.05 were considered to be statistically significant.

## Supporting information

Video S1. A pseudovirus-producing 293FT induces syncytia formation in 293T-hACE2 cells.

## Supplementary Data

Supplementary data is available online.

## Acknowledgement

This work was supported by Ministry of Science and Technology of China (2019YFA0802103, 2021ZD0203900); Department of Science and Technology of Zhejiang Province (2021C03104); Guangzhou Science Innovation and Development Program (201803010092); Shenzhen-Hong Kong Institute of Brain Science (NYKFKT2019006); the National Natural Science Foundation of China (92253302).

