## Supplementary Figure S1 for "A pseudovirus-based method to dynamically mimic SARS-CoV-2-associated cell-to-cell fusion and transmission"

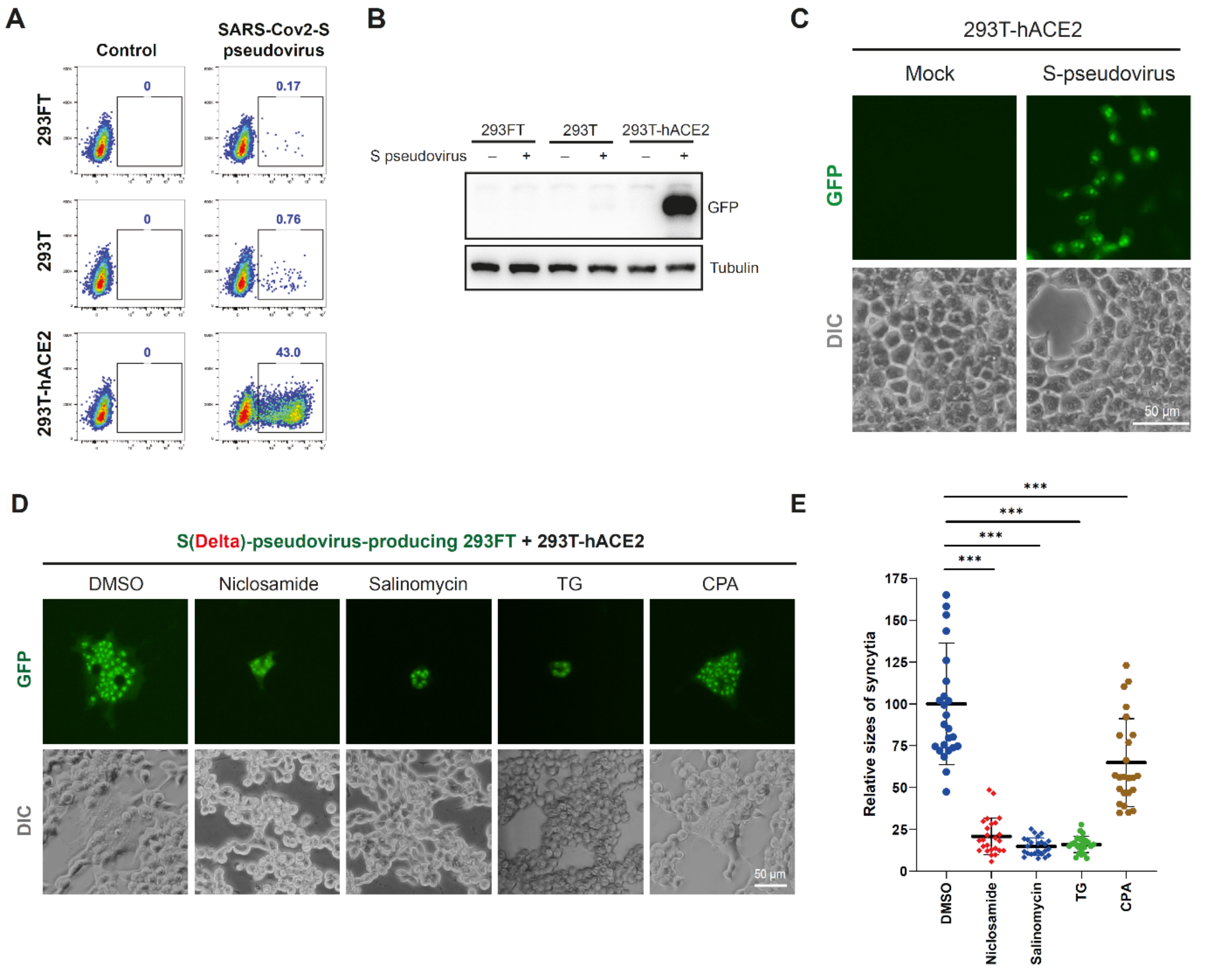


**Figure S1. 293FT generating S pseudovirus can induce cell fusion in hACE2-positive 293T cells.**

**(A** and **B)** SARS-CoV-2 S pseudovirus can effectively infect 293T-hACE2, but not 293T or 293FT cells. 293T, 293FT or 293T-hACE2 cells were incubated with 1 mL DMEM containing S pseudovirion, and seeded in 12-well plates. 24 h later, culture media were changed to fresh DMEM. Cells were cultured for another 24 h. Infection efficiencies were analyzed by flow cytometry (A) or anti-GFP western blot analysis (B).

**(C)** S pseudovirus infection in 293T-hACE2 cells cannot induce syncytia formation. 293T-hACE2 cells was infected by S pseudovirus supernatant. After 48 h, images were taken by Olympus IX73. Scale, 50 μm.

**(D** and **E)** This pseudovirus-based method can estimate the inhibitory effects of drugs on Delta S-mediated cell fusion. DMSO, Niclosamide (1μM), Salinomycin (1μM), thapsigargin (TG) (1μM) or cyclopiazonic acid (CPA) (5μM) was added to the mixture of 293T-hACE2 cells and 293FT cells producing Delta S pseudovirus. Images were shown in (D), and the quantitative analysis of the syncytia sizes is in (E). Scale, 50 μm. Data shown are mean ± SD, ****P* < 0.001. (Student's *t*-test, n = 20).

**Video S1. A pseudovirus-producing 293FT induces syncytia formation in 293T-hACE2 cells.** A single 293FT cell generating S pseudovirus was mixed with 293T-hACE2 cells. Cells were seeded in 29 mm dish and monitored by the Olympus SpinSR real-time live cell imaging system. Cell images were acquired every 5 min for total 15 h and then combined to generate a video.
